# Principal Components Analysis fails to recover phylogenetic structure in hominins

**DOI:** 10.1101/2025.10.31.685754

**Authors:** Levi Y. Raskin, Maja Šešelj, Bárbara D. Bitarello, Sofia Stroustrup, Jacky K. Li, John Huelsenbeck

## Abstract

**Objectives:** Paleoanthropologists often utilize geometric morphometrics and principal components analysis (PCA) to interpret shape variation within the hominin fossil record. It is common practice to interpret proximity in principal components (PC) space among taxa as indicative of not just morphological, but also phylogenetic affinity. This interpretation, however, has not been directly evaluated for hominins.

**Materials and Methods:** First, we inferred the posterior distribution of hominin phylogenetic trees and subsampled trees from this distribution. On these phylogenies, we simulated 2D and 3D geometric morphometric datasets and traditional morphological datasets, containing traits analogous to measurements of size or length, with varying numbers of landmarks or traits and evolutionary rates. On each dataset, we conducted a PCA and used neighbor-joining to infer evolutionary relationships from the PC scores of each taxon. We measure the difference between the PCA tree and sampled tree with subtree pruning and regrafting distance and Robinson-Foulds distance.

**Results:** PCA trees inferred from traditional morphometric data were identical to the sampled tree in 0.11% of datasets when we only considered PC axes 1 and 2, and in 2.9% of datasets when we considered all axes. No PCA tree inferred from any of the 2,400,000 shape datasets was identical to the sampled tree, regardless of the number of axes.

**Discussion:** Phylogenetic interpretations of the hominin fossil record based on proximity in PC space are inherently flawed and likely to be erroneous. Arguments in the hominin systematics literature based on PCA should therefore be reevaluated using phylogenetically-informed alternatives.

## Introduction

Principal Components Analysis (PCA) is a widely applied statistical technique. It reduces multivariate data into sets of orthogonally constrained vectors that explain decreasing proportions of the dataset’s variation (McVean, 2009; Rohlf, 1990). Along with related multivariate statistical methods (such as canonical variate analysis, CVA; Rohlf, 1990), it is especially useful for depicting complex morphological variation and reducing shape space to an easily visualized two- or three-dimensional plot (e.g., Hautavoine et al., 2024). These methods have long been considered the key to interpreting geometric morphometric (GM) datasets (e.g., Rohlf, 1970, 1971), where PCA is a common next step following landmark acquisition and Procrustes analysis, which removes variation in specimen orientation and differences in size. It is unclear, however, to what extent PCA is a predictive tool for phylogenetic relationships rather than simply an exploratory analysis. By design, the method identifies vectors that explain decreasing proportions of variance that must be *mathematically* orthogonal to each other (i.e., independent), ignoring all non-orthogonal vectors even if they were to explain the next sequential proportion of variance. As a result, principal components (PCs) axes identified by a PCA are not necessarily the most informative *biologically* (Cardini et al., 2019).

Nevertheless, paleoanthropologists frequently interpret proximity in PC space as indicative of morphological and thus evolutionary affinity or relatedness (e.g., Davies et al., 2019, 2020, 2024; Dykes, 2016; Gómez-Robles et al., 2007; Harvati, 2003; Harvati et al., 2019; Hautavoine et al., 2024; Holliday et al., 2010; Lague, 2014; Lague et al., 2008; Liu et al., 2022; Mori et al., 2020; Neves et al., 2024; Pan et al., 2016, 2020; Richmond & Jungers, 2008; Skinner et al., 2008; Zanolli et al., 2025). Much of the current knowledge regarding hominin diversity, relationships, and variation comes from studies that use PCA as the sole or primary statistical method. For example, Beaudet et al. (2019) conducted a 3D geometric morphometric (3DGM) study of the bony labyrinth of the South African hominin StW 573 and, based on the PCA, suggested that this fossil should be considered evidence of the ancestral morphotype for South African *Australopithecus*. Hershkovitz et al. (2021) went further and explicitly used PCA to inform their systematics of a newly discovered Middle Pleistocene (MP) hominin fossil from central Israel. Using 3DGM and PCA, Hershkovitz et al. (2021) constructed a neighbor-joining tree between the mean shapes of various prior-specified *Homo* groups and the newly discovered fossils. From the PCA plots and these neighbor-joining trees, Hershkovitz et al. (2021) concluded that these fossils represent the source population to MP European hominins. Hershkovitz et al. (2021) explicitly treated PC-space as a direct proxy for phylogenetic relatedness, an assumption that has yet to be rigorously tested.

The assumption that PC space reflects evolutionary relationships has been challenged over the years (e.g., Adams et al., 2011; Bookstein, 2019; Cardini et al., 2019; Rohlf, 2021). Possible alternatives, such as the phylomorphospace analysis (PA, Rohlf, 2002), phylogenetic principal components analysis (Phy-PCA, Revell, 2009) or the phylogenetically aligned principal components analysis (PACA, Collyer & Adams, 2021), however, have not been widely incorporated in paleoanthropology. Most recently, Mohseni & Elhaik (2024) argued that a PCA of a 3DGM dataset is unable to reproduce known papionin genetic relationships. While this work provides evidence that PCA is inappropriate for systematics, a direct test in a hominin context is still lacking. Here, we elaborate on Mohseni & Elhaik (2024) by directly testing the assumption that PC-space recovers phylogenetic structure in hominins via phylogenetic simulation.

## Materials and Methods

### Simulations

We used phylogenetic simulation to test whether distance in PC space can accurately recover hominin topologies. Phylogenetic simulation is commonly used to evaluate statistical methods in phylogenetics as it allows known generative parameters, such as the known tree topology, to be contrasted against an inferred tree (e.g., Huelsenbeck, 1995). We started by estimating the posterior distribution of the phylogenetic tree topology with the Mongle et al. (2023) character matrix and the Markov k-state (Mk) model (Lewis, 2001). Unlike Mongle et al. (2023), we treated all characters as unordered, as this represents the most general treatment of the character matrix and avoids imparting prior beliefs about the evolutionary history of the coded anatomy, which may change with more research. This posterior distribution accounted for the uncertainty in both the hominin phylogenetic tree topology (how the taxa relate to one another) and the branch lengths of the hominin tree (how much evolutionary change accumulates on each branch). Thus, our simulated trait data were conditioned on the uncertainty in the inferred hominin tree. We sampled 1,000 post-burn-in trees from the posterior distribution to serve as our “true” phylogenetic trees. This workflow is shown schematically in Figure 1.1-1.3.

**Figure 1:**
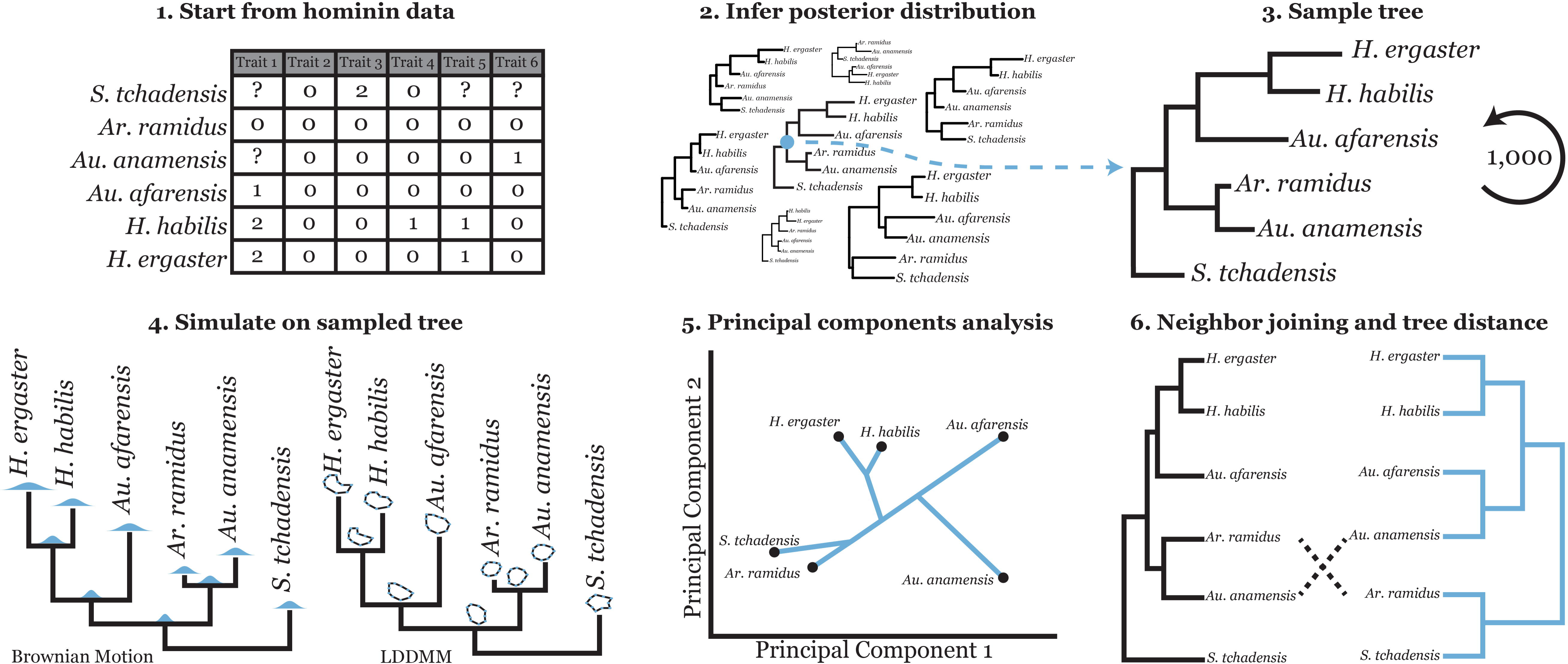
simulation and analysis workflow. (1) Starting with the Mongle et al. (2023) character matrix (21 OTUs, 6 of which shown here), (2) we infer the posterior distribution of phylogenetic tree topology from the character matrix. The posterior probability of each tree topology is represented by the size of the tree, with larger trees indicating a greater posterior probability. (3) We sample a tree from the inferred posterior distribution, referred to as the “true” tree. (4) Then, we simulate two types of data: traditional morphological traits under Brownian Motion and shape traits measured by GM under Large Deformation Diffeomorphic Metric Mapping on the sampled true tree. (5) A PCA analysis is run for each set of traits. (6) A distance matrix is calculated based on PC scores and neighbor-joining is used to infer a “PCA tree”, which is then compared against the true tree. Steps 3-6 are repeated 1,000 times to generate sampling distributions.

First, we simulated both 2D- and 3DGM datasets (Figure 1.4) using the stochastic shape model described in Stroustrup et al. (2025). The stochastic shape model is based on the work by Kunita (1990) and inspired by the large deformation diffeomorphic metric mapping framework (LDDMM; Arnaudon et al., 2019; Younes, 2010). Here, for simplicity, we refer to this stochastic shape model as “LDDMM”. LDDMM models the evolution of landmarks while ensuring they do not cross one another. This model has two parameters: *⍺*, the rate of change, and *σ*, how integrated landmarks are to one another. We simulated datasets with 10, 25, and 50 2D and 3D landmarks, for *⍺* = 0.1, 0.2, 0.3, and 0.4. While *⍺* is straightforward in the LDDMM model (larger *⍺* results in more change over the same period of time), *σ* is less intuitive. High *σ* means that landmarks operate increasingly as a unit wherein a slight change in one landmark affects the position of all the other landmarks, whereas low *σ* results in landmarks evolving increasingly independently from each other, with a small change to one landmark only affecting the position of that landmark. Like *⍺*, *σ* is mathematically constrained to be positive. We fixed *σ* to 1.0 for all values of *⍺* as the interaction between *⍺* and *σ* may result in an equifinality in our shapes: a low *⍺* and low *σ* could result in similar shapes to a high *⍺* and a medium *σ*. Fixing *σ* allowed us to mitigate this potential confounder, while still simulating reasonable, realistic shapes.

We chose these dataset sizes to simulate typical 2D and 3D geometric morphometric datasets in paleoanthropology (e.g., Gómez-Robles et al., 2007; Harvati & Hublin, 2012). Any study using semilandmarks and mesh data itself may have many more landmarks. However, due to computational limitations, we were restricted to sizes that would be typical of datasets incorporating type I and II landmarks. We started all simulations from either a circle or a sphere with evenly distributed landmarks. We did not incorporate conflicting phylogenetic signals into these simulations. For each unique modeling condition (number of dimensions, number of landmarks, and *⍺*), we simulated 100 datasets, resulting in 2,400 unique datasets simulated on each of the 1,000 sampled trees.

Second, we simulated uncorrelated traditional morphological data. These data can be considered analogous to linear measurements in studies like Neves et al. (2024) and Cooper & von Cramon-Taubadel (2025). We simulated datasets under Brownian Motion with 10, 25, 50, 100, 250, and 500 characters. BM is a powerful and common model used to simulate traditional morphological traits and is the most simplistic model for character evolution (see Soul & Wright, 2021). We chose these character matrix sizes to simulate a range of possible datasets, from relatively small to unusually large datasets. For each sampled dataset size, we populated the dataset with traits that evolve at a rate of 0.1, 1, or 10. We also simulated datasets with rate variation across traits. In this case, we drew the rate of change for each trait from a gamma distribution with the following parameters: shape = 1, scale = 10 (mean rate of change is 0.1); shape = 1, scale = 1 (mean rate of change is 1); and shape = 10, scale = 1 (mean rate of change is 10). To explore the effect of conflicting phylogenetic signals, we swapped the simulated value for two randomly sampled taxa for 0%, 10%, 25%, and 50% of traits in each dataset. For each combination of parameters (number of characters, rate of evolution, and proportion of traits with a phylogenetic signal conflict), we simulated 25 datasets, resulting in 5,400 unique datasets simulated on each of the 1,000 sampled trees.

Bayesian inference of phylogeny was done in RevBayes (Höhna et al., 2016), LDDMM simulations of geometric morphometric datasets were written in C++, and BM simulation of traditional morphological datasets were done in R version 4.4.1 with the *fastBM* function from the phytools R package (Revell, 2012). Code to reproduce these analyses is available at github.com/Levi-Raskin/PCAPhylogenetics.

### Principal Component Analyses

For each simulated traditional morphological dataset, we conducted a PCA (Figure 1.5) using the *prcomp* function from the stats R package and, using PC scores, constructed a Euclidean distance matrix using the *dist* function from the stats R package. For each simulated morphometric dataset, we ran two analyses. In the first approach, we used the raw shape data (referred to as “unaligned”) and the *prcomp* function to conduct the PCA. In the second approach (referred to as “aligned”), we used the geomorph R package (Adams & Otárola-Castillo, 2013) function *gpagen* to align, rotate, and scale the simulated shapes with a Generalized Procrustes Analysis before conducting a PCA with *gm.prcomp*.

### Distances and Tree Inference

For each dataset, we calculated two Euclidean distance matrices: first, from principal component axes one and two (PC1, PC2), and second, using scores from all PC axes. From the distance matrix, we used the *nj* function from the ape R package (Paradis & Schliep, 2019) to calculate the neighbor-joining phylogenetic tree (referred to throughout as “PCA tree”) from the distance matrix (Figure 1.6). We chose neighbor-joining, which minimizes distance between taxa (Saitou & Nei, 1987), as it is a consistent and repeatable analogue to the implicit and explicit usage of PCA for systematics. We then calculated the Robinson-Foulds distance (RF distance, Robinson & Foulds, 1981) between the PCA tree and the “true” phylogenetic tree using *dist.topo* (Paradis & Schliep, 2019) and the exact number of subtree pruning and regrafting (SPR) moves needed to transform the PCA tree into the true tree (SPR distance, using rSPR, Whidden et al., 2014). These analyses calculated what is effectively the topological error between the true tree and the tree inferred from PC space. Here, we ignored branch length differences, as we are concerned with the utility of PCA for systematics and not with the utility of PCA to estimate evolutionary time between taxa. To contextualize these differences, we constructed a null distribution by generating 10,000 pairs of random trees with 21 tips using the *rtree* function (Paradis & Schliep, 2019) and calculating the RF and SPR distances between the two trees. This null distribution allows us to test whether the distance between a PCA tree and a true tree is smaller than expected if they were truly a random pair of trees.

## Results

### Simulating GM data

Our goal was to explore whether PCA is able to recover underlying phylogenetic structure; we found that it does not. By simulating 2D shapes, 3D shapes, and traditional morphological datasets, we were able to directly assess the relationship between observed data at the tips of a phylogeny and the known true phylogeny, a luxury impossible with real-world data. To start, we ensured that our shape simulations produced landmark configurations across rate conditions comparable to hominin anatomy. Figure 2 showcases arbitrarily selected 2D and 3D shapes, along with the phylogenetic tree on which they were simulated. At low rates of evolution (e.g., *⍺* = 0.1), the 2D shapes shown here resembled circles (the starting shape at the root), as would be expected. These low-rate shapes are intended to be comparable in shape and diversity to relatively conserved structures, such as molar occlusal outlines or long bone cross-sectional shapes. At high rates (e.g., *⍺* = 0.4), the simulated shapes shown here were divergent from the original circular shape and model structures for which there is substantial morphological diversity across the hominin clade, such as the shape of the mandibular symphysis. The 3D shapes likewise approximated a range of real-world morphologies. At lower rates, the simulated shapes at the tips were roughly spherical (the original shape at the root), approximating structures like the femoral head. At medium rates (e.g., *⍺* = 0.3), we could model 3D structures that show some variation across the clade but are relatively more constrained, such as the calvaria or neurocranial morphologies. At higher rates, there was significant diversity across the clade, modeling real-world 3D structures that are highly variable, such as sinus morphology. While the shapes shown in Figure 2 are isolated examples and not necessarily indicative of the entirety of the simulated datasets, they both match our expectations of the amount of diversity for each rate and are representative of what is typical within the simulated datasets. Here, we highlight these specific examples to demonstrate that our simulation framework can produce simulated data that are both consistent with statistical expectations and biological expectations.

**Figure 2:**
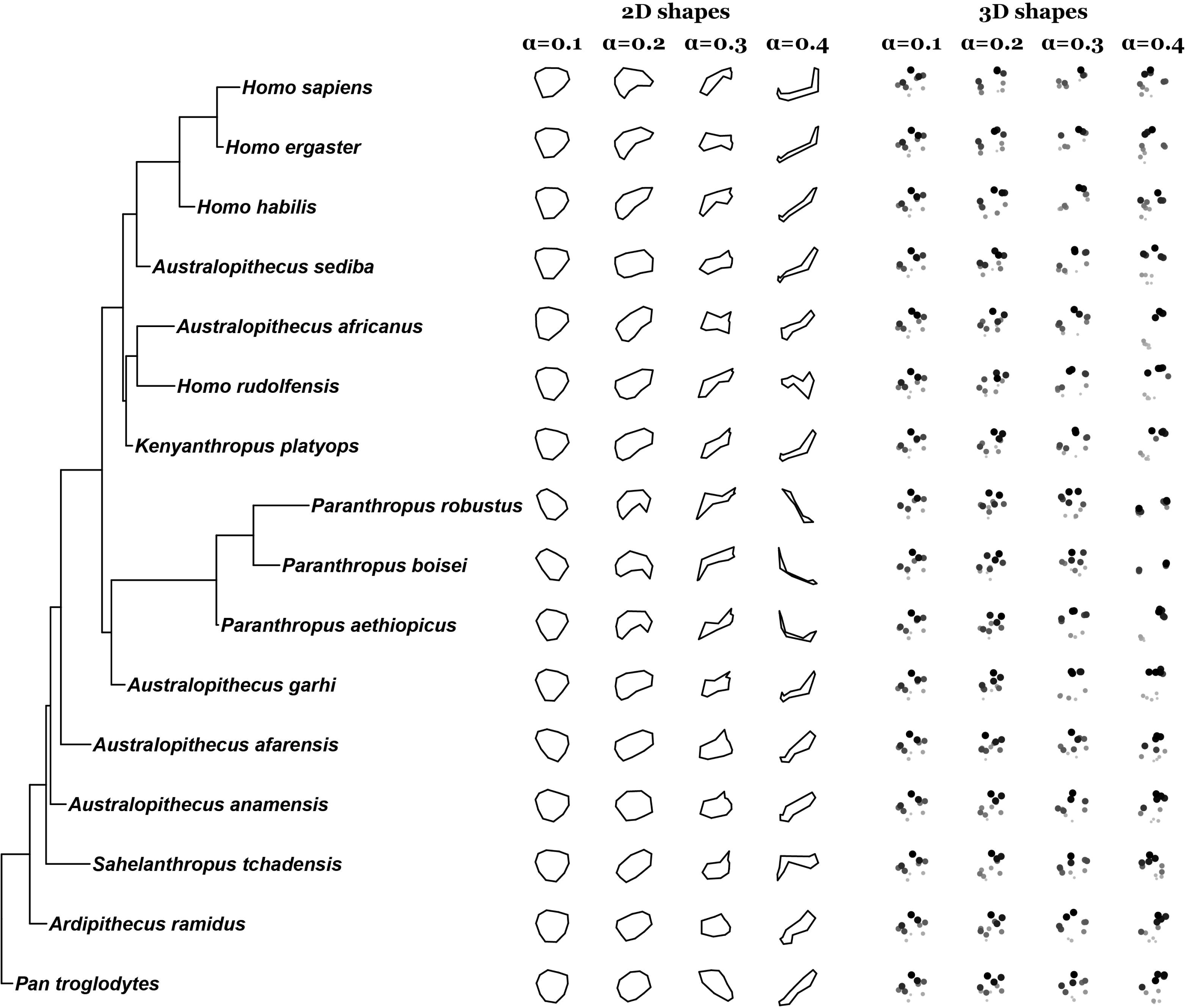
example simulated shapes across rates. Left, the hominin and *Pan* subtree from an example sampled phylogenetic tree. Center, simulated 2D shapes across the four rate conditions from four independent simulations. Landmarks are plotted as points with neighboring landmarks connected with straight lines. Right, simulated three-dimensional shapes across the four rate conditions from four independent simulations. Landmarks are shown as points, with edges depicting the convex hull of the configuration. The X and Y coordinates are plotted along the horizontal and vertical axes, respectively; the Z coordinate is plotted with point size and shading intensity with closer landmarks depicted larger and darker.

We note that simulating under the LDDMM model produced shapes recognizable to an anatomist and produced synapomorphies recognizable to a systematist. For example, the 2D shapes simulated with *⍺* = 0.2 depicted in Figure 2 for the *Paranthropus* clade all have a concave notch at the bottom of the shape. In this simulation, the notch most likely evolved once and represents a shared, derived feature of the *Paranthropus* clade. Interestingly, independently (i.e., a homoplasy), *Homo sapiens* and *H. ergaster* both present a similar notch. This model could also reproduce the loss of anatomical structures over time. For example, for the 2D shapes simulated with *⍺* = 0.3 depicted in Figure 2, *Sahelanthropus tchadensis*, *Australopithecus anamensis*, and *Au. afarensis* all have a triangular protrusion on the top of the shape, which is lost in later hominins.

### PCA fails to recover phylogeny from GM data

Despite these desirably realistic aspects of the LDDMM model, PCA was unable to recover the underlying phylogenetic structure in the GM data. We did not observe a single tree based on PC1 and PC2 or all PCs, across all rates, number of landmarks analyzed, and dimension of landmarks, that was identical to the true tree on which the data was simulated for both the unaligned and the aligned GM datasets (Figure 3).

**Figure 3:**
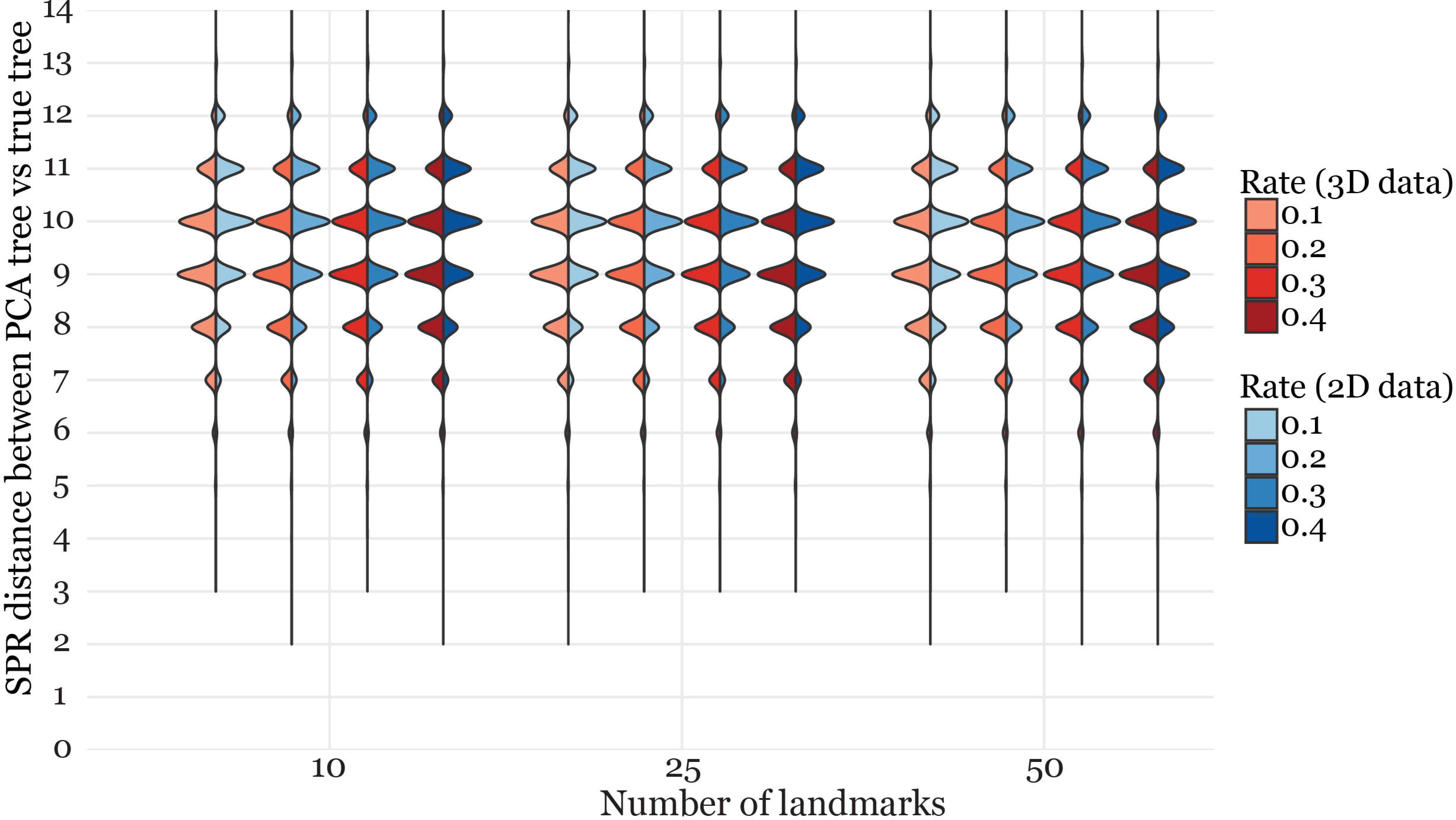
SPR distances for unaligned geometric morphometric PCA trees inferred from PC1 and PC2. SPR distances were calculated between the associated true tree and PCA trees inferred from datasets with varying numbers of landmarks for both 2D and 3D data with evolutionary rates fixed at (from left to right) 0.1, 0.2, 0.3, and 0.4.

For our unaligned GM data, the median SPR distance across all parameters was 10 (*p* = *0*.*057*, calculated against our null distribution), the minimum was 2, and the maximum was 14. The median RF distance was 32 (*p* = *0*.*027*), the minimum was 6, and the maximum was 36, the largest possible RF distance for trees with 21 tips (Kuhner & Yamato, 2015). Using all PCs similarly resulted in distant trees, though again we did not observe a single PCA tree identical to the true tree. The median SPR distance across all parameters was 9 (*p* = *0*.*003*), the minimum observed was 2, and the maximum was 14. The median RF distance was 32 (*p* = *0*.*027*), the minimum observed was 4, and the maximum observed was 36.

For the aligned GM data, the median SPR distance across all parameters was 11 (*p* = *0*.*556*), with a minimum of 3, and maximum of 14. The median and maximum RF distances were 36 and the minimum was 8. Across all PCs, the median SPR was 10 (*p* = *0*.*057*), the minimum was 2, and the maximum was 14. The median RF distance was 34 (*p* = *0*.*210*), the minimum was 8, and the maximum was 36.

### Increasing number of landmarks reduces PCA tree distance

We investigated whether increasing the number of landmarks affected the distance between the PCA tree and the true tree. For the unaligned data, we found that for all analyses and parameters, every increase in the number of landmarks resulted in a significantly reduced distance between the PCA tree and the true tree (*p* ≤ *1*.*97* ∗ *10*^−*11*^, Mann-Whitney U test [MWU], SOM table S1). However, the magnitude of the shift in means in every case was <1 RF or SPR distance. Similarly, while 3D landmarks perform significantly better than 2D landmarks for all PC analyses across all parameters for both SPR and RF distances, the magnitude of the shift in means was small, with the largest shift a mean decrease in RF distances by 2.12 from the analysis of all PCs.

The effects of increasing the number of landmarks on SPR were comparable for our aligned dataset. Every increase in the number of landmarks for both the PC1 and PC2 and the full PC analysis resulted in a significantly reduced SPR distance between the PCA tree and the true tree (*p* ≤ *2*.*2* ∗ *10*^−*16*^, Mann-Whitney U test [MWU], SOM table S1), with the magnitude of the shift in means always <1 SPR distance. An exception was the comparison between trees using 10 and 25 landmarks, where for PC1 and PC2 we observed a mean RF distance of 34.55 and 34.63, respectively. For both PC1 and PC2 and across all PCs, we observed a significant increase in both RF and SPR distance in PCA trees inferred from aligned 3D relative to 2D landmarks (*p* ≤ *2*.*2* ∗ *10*^−*16*^, Mann-Whitney U test [MWU], SOM table S1). While significant, these shifts were qualitatively minor (<1 RF or SPR distance) in all cases.

### PCA fails to recover phylogeny from traditional morphological data

Simulated traditional morphological datasets, analogous to measurements like linear distances, mass, volumes, and angles, likewise failed to reliably recover phylogenetic structure. For PC1 and PC2, across all parameters, only 3,953 out of the total 3,600,000 PCA trees (0.11%) were identical to the true tree. The median SPR distance across all parameters was 5 (the smallest SPR distance in our null distribution was nine), and the maximum was 13. The median RF distance was 14 (the smallest RF distance in our null distribution was 28) with a maximum of 36. An example PCA, PCA tree, and true tree are depicted in Figure 4. Using all PCs instead of just PC1 and PC2 modestly improved accuracy. Across all parameters, 103,136 out of the total 3,600,000 PCA trees (2.9%) were identical to the true tree. The median SPR distance across all parameters was 4, with, again, a maximum observed SPR distance of 13. The median RF distance was 10, and again, the maximum RF distance was 36.

**Figure 4:**
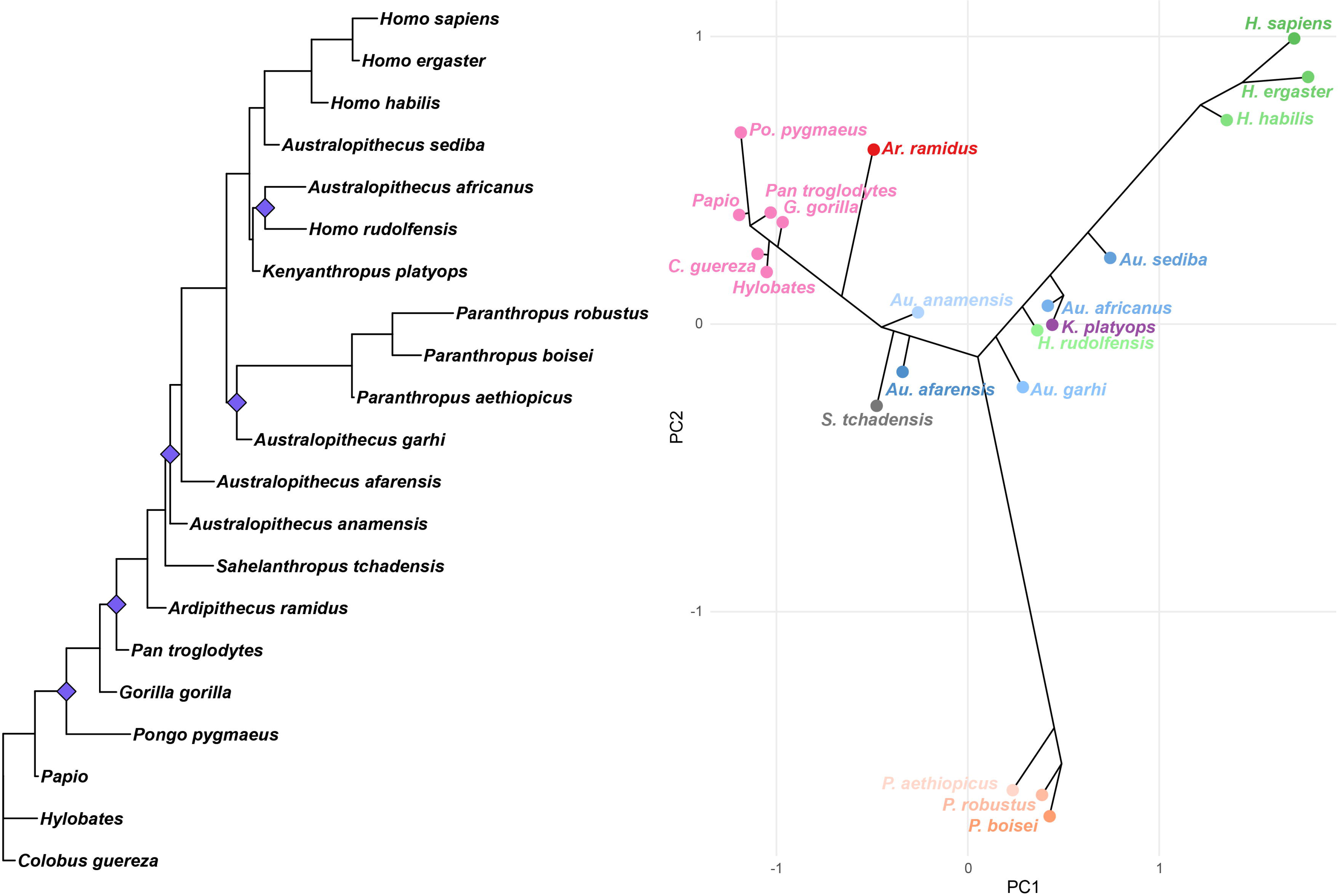
example PCA and neighbor-joining tree estimated from a traditional morphological dataset. Left, an example phylogenetic tree sampled from the posterior distribution inferred using RevBayes. This is the same topology depicted in Figure 2. Right, PCA and neighbor-joining tree (based on PCs 1 and 2) constructed from a 50-trait dataset simulated under Brownian motion on the example tree (left) with a fixed rate of 0.1 and no conflicting phylogenetic signals. Extant non-hominin great apes are colored pink. Other colors group hominin genera, with different shades of each color for species within each genus. The plotted neighbor-joining tree depicts relationships among taxa, but does not accurately represent estimated branch lengths. Purple diamonds on the example tree indicate the subtree pruning locations needed to transform it into the neighbor-joining tree with subtree pruning and regrafting moves.

### Increasing the number of traditional traits reduces PCA tree distance

Increasing the number of characters in the character matrix increased the number of PCA trees identical to the true tree. For PC1 and PC2, we did not observe a single PCA tree identical to the true tree for character matrices with 10 characters (out of 600,000 character matrices simulated for each size, Figure 5). We observed seven PCA trees (0.001%) identical for a character matrix with 25 traits, 50 PCA trees (0.008%) identical for a character matrix with 50 traits, 264 PCA trees (0.044%) identical for a character matrix with 100 traits, 1,175 PCA trees (0.196%) identical for a character matrix of size 250, and 2,457 PCA trees (0.409%) identical for a character matrix of size 500. The pattern remains the same for PCA trees inferred from all PCs. We observed two identical PCA trees (0.0003%) for character matrices of size 10, 172 (0.029%) for character matrices of size 25, 1,896 (0.316%) for character matrices of size 50, 9,785 (1.631%) for character matrices of size 100, 34,292 (5.715%) for a character matrix of size 250, and 56,989 (9.498%) identical PCA trees for a character matrix of size 500.

**Figure 5:**
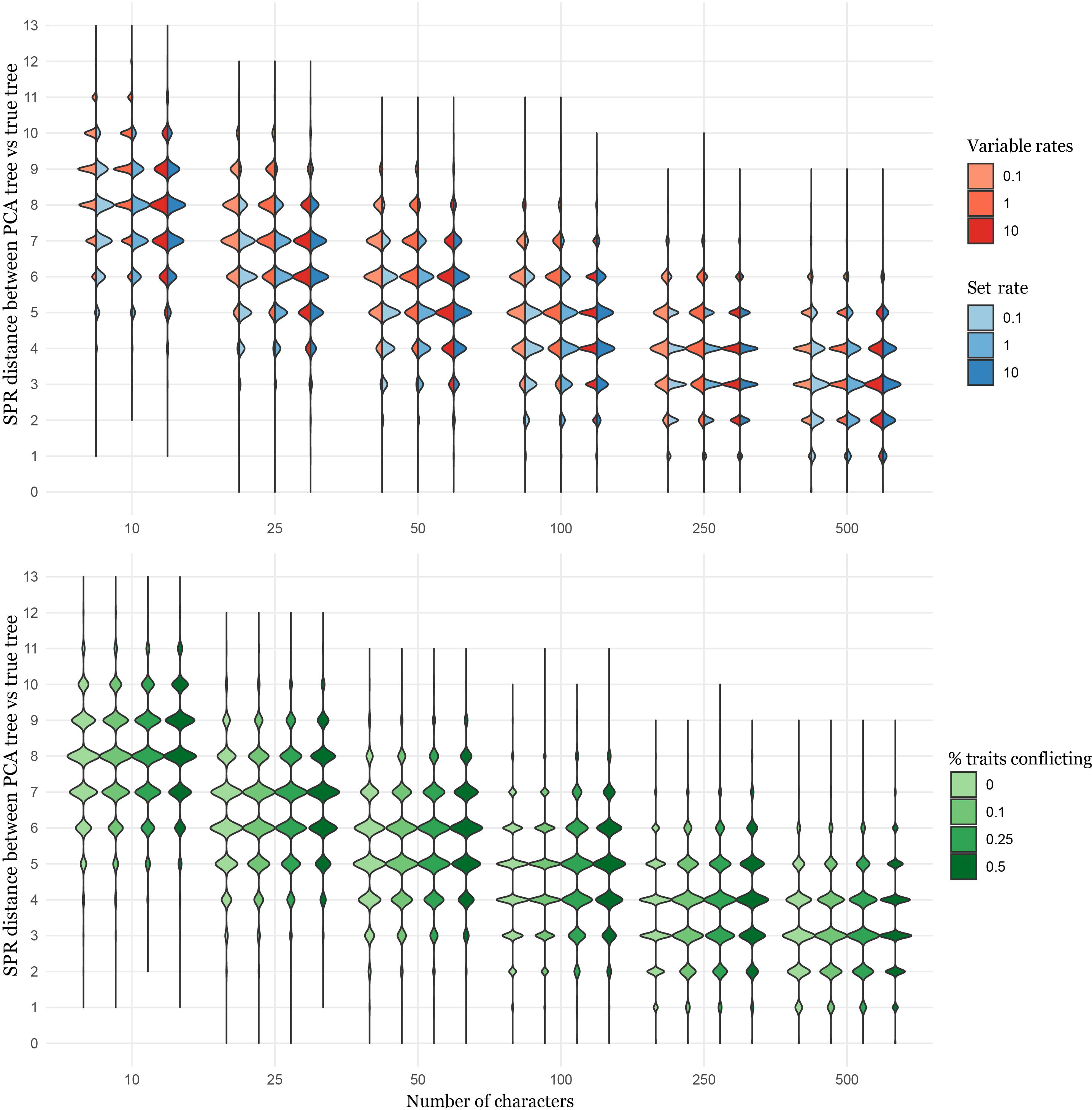
SPR distances for traditional morphological PCA trees inferred from PC1 and PC2 across character matrix size, rates, and proportion of the data with conflicting phylogenetic signals. Top, SPR distances for datasets of varying sizes with evolutionary rates drawn either from a gamma distribution with a mean of (from left to right in each group) 0.1, 1, or 10 or a fixed evolution rate of 0.1, 1, or 10. Bottom, SPR distances for datasets of varying sizes with different proportions of characters presenting conflicting phylogenetic signals.

### Conflicting phylogenetic signals and variable rates of evolution increase PCA tree distance

We investigated the effects of rate variation across characters by comparing the accuracy of PCA-based systematics for character matrices where the rate of change for each trait is drawn from a gamma distribution to character matrices where the rate of change for each trait is set to the expectation of the gamma distribution (i.e., a character matrix where rates of evolution are drawn from a gamma with parameters shape = 1 and scale = 10 compared to a character matrix simulated with a set rate of 0.1). For the PCA trees inferred from both PC1 and PC2 and all PCs, we found that for almost every single size of character matrix without conflicting phylogenetic signals, the variable rates character matrix had a significantly higher SPR distances from the true phylogenetic tree (*p* ≤ *3*.*68* ∗ *10*^−*4*^, Mann-Whitney U test [MWU], SOM table S2). The sole comparison between variable rates and set rate PCA tree SPR distances that was not significant was for datasets with 500 characters, inferred using all PCs, and with an expectation and set rate of 10 (*p* = *0*.*158*, Mann-Whitney U test [MWU], SOM table S2). This same pattern holds for RF distances. Similarly, as the proportion of the character matrix with conflicting phylogenetic signals increases, the mean RF and SPR distances both significantly increase for most character matrix sizes (*⍺* = *0*.*05*, MWU, SOM table S2). The exceptions were all character matrices of size 500, where, in some comparisons (e.g., proportion conflicting = 0 vs. proportion conflicting = 0.1), the mean RF distance actually decreased. However, these decreases were marginal and never statistically significant.

## Discussion

We applied phylogenetic simulation methods and a newly developed model to investigate the utility of PCA for hominin systematics. We found that PCA is unable to accurately recover the underlying phylogenetic structure in both traditional morphological and morphometric datasets, as we discuss below. Because a concerning proportion of the hominin systematics literature relies on an inherently flawed interpretation of PCA, we took great care to ensure that our results are truly generalizable to the hominin fossil record and robust to a wide variety of conditions.

Both types of simulated data (traditional morphological data and GM data) model real paleoanthropological datasets that, in concert with PCA and other multivariate analyses, have directly informed hominin systematics. First, we performed our simulations on 1,000 trees sampled from the posterior distribution of tree topologies. This sampling ensured that the true trees underlying our simulations were as representative as possible of hominin phylogeny (Huelsenbeck et al., 2000). While we have taken care to ensure that our “true” trees are representative, we make no specific claims about the real hominin tree, and refer readers to other studies (e.g., Mongle et al., 2023; Parins-Fukuchi et al., 2019).

However, the above does not reduce our ability to address our research question, and, in fact, we are able to do that by accounting for uncertainty in both topology and branch lengths, conditioned on the Mongle et al. (2023) dataset. The trees we sampled from the posterior distribution have branch lengths that represent the relative amount of evolutionary change between tips. We simulated our datasets on trees with a variety of branch lengths proportional to the probability of that branch length, which accounts for the uncertainty in the evolutionary time separating each taxon. The rate of change for our simulated characters is directly related to the branch lengths of our true trees. Under most evolutionary models, larger branch lengths and smaller rates result in the same expected probability distribution as smaller branch lengths and larger rates (Rannala, 2002). For this reason, we chose our rates to produce a range of reasonable data at the tips across the range of branch lengths in our posterior distribution. While these simulated rates may not match real-world estimates for hominin traits, our chosen rates result in a range of biologically plausible structures. The resultant shapes range from one extreme, where traits are highly conserved (low rates), to the other extreme, where shapes may become biologically implausible (high rates), but covering the range of hypothetical anatomical structures that would be interesting to a paleoanthropologist. Despite conditioning on branch length uncertainty and our careful selection of our model parameters, we observed poor performance of PCA in recovering plausible hominin phylogenies despite PCA’s long-standing application to that question.

As an example, while the differences between the true tree and the PCA tree depicted in Figure 4 may seem minor (16 out of 21 tips correctly positioned for that PCA tree), these differences were still sufficiently large to preclude the usage of PCA in hominin systematics, particularly contrasted with the longstanding usage of BM and similar models to infer phylogenies (Felsenstein, 1973). While Bayesian and maximum likelihood methods are not guaranteed to recover the true phylogenetic tree, they are statistically consistent methods with approaches that can quantify the uncertainty in the recovered topology (e.g., Felsenstein, 1985; Huelsenbeck et al., 2000). PCA is not necessarily statistically consistent for topology, does not have a clear ability to accommodate uncertainty in the estimated trees, and, as we showed here, is not accurate for realistic dataset sizes. Furthermore, PCA is known to be biased in cases where sampling is uneven (McVean, 2009). Our simulated datasets contained no missing data, a luxury not present in the true hominin fossil record (McRae & Wood, 2025), nor present in papers that directly utilize PCA to inform systematic arguments (e.g., Davies et al., 2024; Harvati et al., 2019; Hershkovitz et al., 2021). Thus, our analyses assessed the “best case scenario” for PCA as an inference tool, free of common biases seen in fossil datasets. Similarly, we used NJ to infer the phylogenetic tree, despite it occasionally producing counterintuitive phylogenetic trees. If one was to claim phylogenetic relationships based on the PCA alone, one might conclude, for example, that the central cluster we observe in the PCA shown in Figure 4 (australopiths, *H. rudolfensis*, and *S. tchadensis*) forms clade. This would signify a larger distance from the true topology than the actual NJ tree inferred based on the PCA (tree overlaid over PCA in Figure 4). In other words, NJ may be modestly more accurate at recovering the underlying phylogenetic structure assumed present in the PC scores than a heuristic, manual approach.

Interestingly, the median PCA tree distance was smaller than our random null distribution in almost every single analysis, with the sole exception of the SPR distances from our unaligned GM data (*p* = *0*.*057*). For our traditional morphological data, the median PCA tree RF and SPR distances were vastly smaller than the smallest distance we observed in our null distribution. Much of this can be attributed to the size of tree space: even though we sampled 20,000 total random trees in the calculation of our null distributions, they represent an infinitesimally small percentage of the entirety of the tree space. Considering how little tree space we sampled for our null distribution, the fact that we see in our null distribution some comparable distances between random pairs of trees and the distances between our true trees and GM PCA trees is a strong indicator that, at least for GM data, PCA is a relatively minor improvement over randomly selecting a tree.

Increasing the number or dimensionality of landmarks produces only a minor reduction in PCA tree distances. Theoretically, this minor effect is expected: contrary to traditional morphological data, where each additional character is an independent datum, a single configuration of landmarks represents a single datum, regardless of the number of landmarks (e.g., Zelditch, 2004). Furthermore, it has been well-established that a high number of landmarks, which is especially the case for datasets that include semilandmarks (as outlines or surface meshes), relative to the number of specimens or groups included in the analyses (i.e., high p/n, Bookstein, 2019) can lead to a distortion of the appearance of plots, inflating between-group variation, and even creating spurious clusters in between-group PCA, or bgPCA (Cardini et al., 2019; Rohlf, 2021). Our traditional morphological data PCA tree distances were smaller than expected of two randomly selected trees, showing a decided improvement over randomness, but in no scenario should the PCA tree be expected to be accurate to the true underlying phylogenetic tree.

Finally, within GM analyses, the unaligned approach outperforms the Procrustes-aligned approach. For real data, the relative rotation of each landmark is not informative. For example, in real-world data, the configuration of landmarks is not consistently rotated relative to the centroid for all three OTUs within *Paranthropus*, but instead an artifact of the data collection process itself. For our unaligned dataset, the entire landmark configuration for the *Paranthropus* OTUs could be rotated relative to the centroid simply because along the branch leading to *Paranthropus*, the ancestral configuration of landmarks was stochastically rotated. Simulated GM data containing more information than their real-world analogue is an artifact of simulations and should be considered in future simulation-based studies of GM data. Procrustes superimposition also mathematically adjusts specimens for size, so our result may also indicate that some proportion of phylogenetically useful information is related to size, resulting in the slightly better performance of unaligned vs. aligned GM data.

Ultimately, the reasons why PCA is unable to recover true phylogenies, especially using GM data, are unclear at this time, but are likely multifactorial. In addition to issues noted above, the PCA itself is not inherently a phylogenetic method, but a way to simplify and visualize complex shape variation by reducing multidimensionality and projecting the data onto axes that are not necessarily the most informative biologically (Cardini et al., 2019). PCA is essentially a phenetic method in absence of phylogenetic data (Adams et al., 2011). In a study using genomic data from two human populations (McVean, 2009), PC1 was able to recover coalescence dates for those two populations, but the results did not hold for three or more populations, suggesting PCA does not lend itself readily for phylogenetic applications. Morphological analyses are even more challenging in that morphological datasets generally consist of a mix of traits, or landmarks, that may or may not be phylogenetically informative or useful, for various reasons (Raskin et al., 2024). Therefore, while a pattern of variation visualized along PC axes might reflect morphological affinity, albeit limited to morphological variance along mathematically (not biologically) informative axes, it does not follow that it also reflects phylogenetic affinity. As Mohseni & Elhaik (2024) note, manual selection of specific PC axes to consider can create the appearance of recovering biological and phylogenetic truth in the PCA, but, when this biological and phylogenetic truth are unrecoverable, picking and choosing PC axes to match assumptions about a group’s biology is insufficient evidence. In our study, there may exist PC axes that, in various combinations, do perfectly recover the true tree, but in practice, these idealized axes are unrecoverable.

Several researchers have proposed alternatives that incorporate phylogenetic information, such as phylomorphospace analysis (Rohlf, 2002) and the similar cladistic morphospace analysis (Stone, 2003), as well as modifications of PCA such as phylogenetic principal components analysis (Phy-PCA, Revell, 2009, but see Polly et al., 2013; Uyeda et al., 2015) and the phylogenetically aligned principal components analysis (PACA, Collyer & Adams, 2021). Possible explanations and solutions for the failure of bgPCA were also recently put forth (Bookstein, 2019; Cardini & Polly, 2020; Rohlf, 2021). More research is needed to further test existing phylogenetically-informed approaches to geometric morphometric analyses, as well as to develop new methods that would allow direct inference of phylogeny from geometric morphometric data.

## Conclusion

PCA is widely used throughout the life sciences in part because it provides an intuitive way to look at multivariate data in a dimension-reduced manner — thus it applies to a variety of biological datasets — that is graphically appealing and has a relatively low barrier to entry. Nevertheless, due to being nonparametric, it can be challenging to link PCA to underlying processes (McVean, 2009). As such, it is in most cases an exploratory tool to uncover axes of variation in data, though in some cases the relationships between underlying process and PC projection are well understood (e.g., the underlying genealogical history of samples and genotype-based PCAs, see McVean (2009). While it is seemingly a useful tool to explore morphological variation and behavior (e.g., Burns et al., 2024; Green & Alemseged, 2012), it is not suitable for morphological systematics, neither explicit (via PCA-informed tree inference, as we have done) nor implied, as others have also argued (Adams et al., 2011; Cardini et al., 2019; Mohseni & Elhaik, 2024).

Using a wide variety of parameters including variation rates and numbers of rates, we showed that, for traditional morphological datasets, PCA is expected to recover the underlying phylogenetic structure for only between 0.11 and 2.9% of datasets. This poor performance is likely an overestimate compared to analyses with real data, given that our simulated datasets had no biases or missing data common in paleoanthropological datasets. Morphometric datasets perform even worse for PCA-based systematics, as we did not observe a single instance where the PCA tree was identical to the true topology in both our Procrustes-aligned and unaligned datasets, and the mean discrepancy between the true and GM PCA-inferred trees was not unusual amongst pairs of random trees.

We expect that these results are robust to the uncertainty in the hominin phylogenetic tree and directly generalizable not just to the hominin fossil record, but paleontology more broadly. Among the instances where the PCA-based trees did recover the known underlying phylogenetic tree, we did not observe any patterns in the generative parameters, except that larger character matrices and less conflicting phylogenetic signals both modestly increase the probability that the PCA-based tree is accurate to the true topology. Despite its widespread use in paleoanthropology, PCA is fundamentally ill-suited and unreliable for inferring hominin evolutionary relationships from morphological data. Studies of hominin systematics that incorporate, and especially those that rely on, PCA as a means of inferring morphological and phylogenetic affinity should therefore be revisited.

## Funding acknowledgment

This material is based upon work supported by the U.S. Department of Energy, Office of Science, Office of Advanced Scientific Computing Research, under Award Number DE-SC0026073 to LYR. This work was supported by a research grant (VIL40582) from Villum Fonden.

